# EvoAug-TF: Extending evolution-inspired data augmentations for genomic deep learning to TensorFlow

**DOI:** 10.1101/2024.01.17.575961

**Authors:** Yiyang Yu, Shivani Muthukumar, Peter K Koo

## Abstract

Deep neural networks (DNNs) have been widely applied to predict the molecular functions of regulatory regions in the non-coding genome. DNNs are data hungry and thus require many training examples to fit data well. However, functional genomics experiments typically generate limited amounts of data, constrained by the activity levels of the molecular function under study inside the cell. Recently, EvoAug was introduced to train a genomic DNN with evolution-inspired augmentations. EvoAug-trained DNNs have demonstrated improved generalization and interpretability with attribution analysis. However, EvoAug only supports PyTorch-based models, which limits its applications to a broad class of genomic DNNs based in TensorFlow. Here, we extend EvoAug’s functionality to TensorFlow in a new package we call EvoAug-TF. Through a systematic benchmark, we find that EvoAug-TF yields comparable performance with the original EvoAug package.

**Availability:** EvoAug-TF is freely available for users and is distributed under an open-source MIT license. Researchers can access the open-source code on GitHub (https://github.com/p-koo/evoaug-tf). The pre-compiled package is provided via PyPI (https://pypi.org/project/evoaug-tf) with in-depth documentation on ReadTheDocs (https://evoaug-tf.readthedocs.io). The scripts for reproducing the results are available at (https://github.com/p-koo/evoaug-tf_analysis).

## Introduction

Deep neural networks (DNNs) have emerged as a promising tool for supervised learning of regulatory genomics data, taking DNA sequences as input and predicting the readouts of functional genomics experiments^1,2^. Due to their overparameterization, DNNs are data hungry, requiring large amounts of data to learn discriminative features that ensure good generalization^3^. However, most functional genomics experiments only observe a limited number of molecular interactions within the context of a cell. For instance, the number of binding sites available for a transcription factor can be limited to the finite number of accessible DNA within a given cell type. Hence, dataset size is a major limiting factor when analyzing functional genomics data with DNNs.

Data augmentation is a widely practiced strategy in machine learning to provide additional training samples. In practice, random transformations that maintain the same training labels are imposed on the input data, leveraging the natural symmetries in the data. For example, in natural images, the objects can undergo affine transformations, flips, color perturbations, and blurs, which do not alter the object’s label. In genomics, the available transformations that can maintain the same training labels are limited to reverse complements and small random shifts^4,5^.

To expand the available augmentations, EvoAug^6^ was recently introduced to provide evolution-inspired data augmentations, including translocations, insertions, deletions, inversions, and mutations. In nature, genetic variation is sampled by evolution to increase phenotypic diversity. Thus, genetic mutations can alter the function of the sequence and, hence, change its label. Nevertheless, EvoAug asserts that the transformed sequences retain the same training label as the wild-type sequence, which can be considered imposing a prior. For instance, small random translocations impose a prior that the activity of motifs is invariant to shifts. Moreover, insertions and deletions impose a prior that the distance between motifs is insignificant. Considered as a prior, evolution-inspired augmentations can introduce a bias in the DNN that may break the underlying rules of *cis*-regulatory grammars in the data. Thus, EvoAug employs a second stage of training that finetunes the DNN on the original, unaltered data, ensuring functional integrity towards the observed biology. This two-stage training curriculum has proven beneficial for genomic DNNs using synthetic data augmentations^6^ and natural data augmentations^7^, as well as for protein-based DNNs^8,9^.

However, the availability of EvoAug has been limited to a PyTorch^10^ implementation, posing a barrier for researchers working in TensorFlow^11^. To address this demand, we present `EvoAug-TF`, a TensorFlow implementation that builds upon the same principles as EvoAug. Below, we describe EvoAug-TF, highlighting its adaptations and unique features that are specific to the TensorFlow framework. Additionally, we present experimental results showcasing the effectiveness of EvoAug-TF in improving the generalization of genomic DNNs.

### Methodology and Implementation

EvoAug-TF adapts the functionality of the PyTorch-based EvoAug framework in TensorFlow (Fig. 1a), including all of the augmentation techniques (e.g., random transversion, insertion, translocation, deletion, mutation, and noise). EvoAug-TF employs the same two-stage training curriculum, where stochastic augmentations are applied online to each mini-batch during training, followed by a finetuning step on the original, unperturbed data. Since EvoAug-TF imposes transformations on the input data while maintaining the same labels as the wildtype, in its current form, EvoAug-TF should be applied to DNNs that output scalars in single-task or multi-task settings. In contrast, profile-based DNNs^5,12,13^ may require transformations to the corresponding labels.

**Figure 1.**
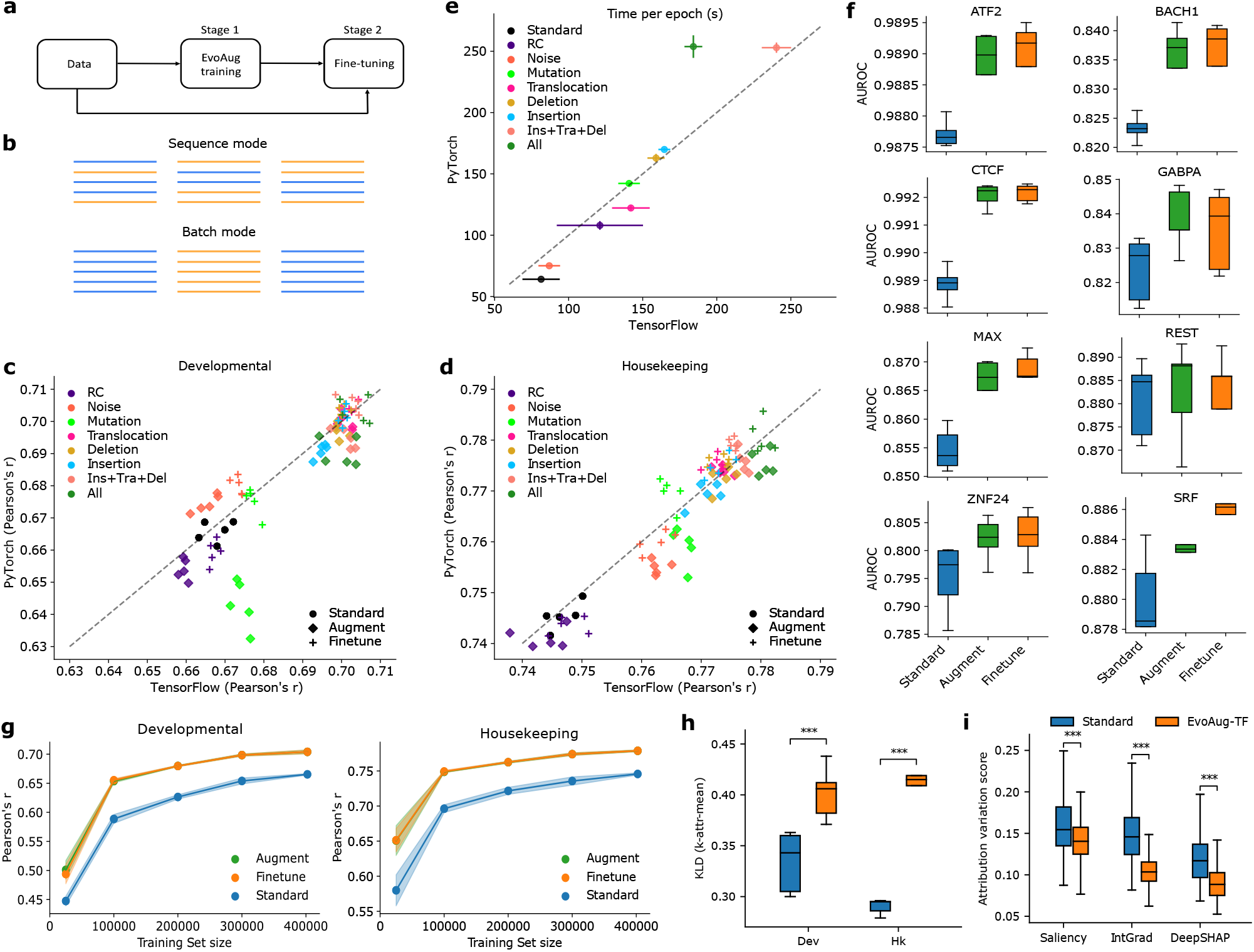
Performance comparison between EvoAug and Standard training. **a** Flowchart illustrating EvoAug-TF’s two-stage training curriculum. **b** Schematic demonstrating the difference between sequence mode and batch mode, where each column represents a mini-batch of sequences and different colors represent different data augmentations. **c, d** Performance comparison between EvoAug (PyTorch-based) and EvoAug-TF (TensorFlow-based) with DeepSTARR models trained with individual or combinations of augmentations (i.e., insertion + translocation + deletion; all augmentations) and finetuned on original STARR-seq data for two promoters: developmental (**c**) and housekeeping (**d**). **e** Comparison of the training time per epoch between EvoAug (PyTorch) and EvoAug-TF (TensorFlow). **c-e** Each scatter plot includes the data from 5 trials with random initialization. **f** Box-plots comparing the performance (area under the receiver operating characteristic curve, AUROC) of a convolutional neural network trained with standard training, EvoAug-TF augmentations in batch mode, and with finetuning on various Chip-seq peak classification tasks. **g** Performance comparison of DeepSTARR models with various training methods fit to training sets with varying levels of down-sampling. EvoAug-TF augmentations employed insertion + translocation + deletion in batch mode. The shaded region represents the standard deviation of the mean across five models with different random initializations. **h** Box-plot comparison of the consistency of patterns within Saliency Maps, as measured by the *k*-attr-mean, for DeepSTARR models with standard training or EvoAug-TF augmentations (All). **i** Box-plot comparison of the attribution variation score, calculated according to the root-mean-squared attribution scores, for attribution maps generated by different methods (i.e., Saliency Maps, Integrated Gradients, and DeepSHAP) using a DeepSTARR model trained with standard training or EvoAug-TF augmentations (All). **h**,**i** Mann-Whitney U test with p-values less than 0.001 (***) and 0.01 (**). Boxes represent five identical models with different random initializations.

Several adaptations were made to EvoAug-TF compared to the PyTorch implementation, incorporating key design choices specific to the TensorFlow version. To ensure compatibility with TensorFlow’s graph mode and optimize training speed, `tf.Tensors` are used as substitutes for NumPy arrays, which were utilized in the PyTorch implementation. Additionally, EvoAug-TF employs `tf.while_loop` to achieve the same effects as the for loops used in the PyTorch implementation. The implementation of EvoAug-TF relies on TensorFlow 2 and its relevant libraries and dependencies; it has been tested thoroughly on versions 2.7 and 2.15.

A key difference between EvoAug-TF and EvoAug lies in the approach to applying augmentations to the mini-batch during training. In the PyTorch implementation, the number of augmentations and the augmentation type are randomly chosen for each sequence. In the TensorFlow implementation, the same number of augmentations and augmentation types are employed for each sequence. However, the selected augmentation type(s) are sampled in a stochastic manner to impose a unique perturbation to each sequence in the mini-batch, similar to EvoAug. This design choice was made to simplify the process of imposing augmentations, while still sampling a high degree of genetic variation. Moreover, for some augmentations – such as translocation, insertion, deletion, and transversion (i.e., reverse-complement) – we offer an additional option to perform the same perturbation across all of the sequences within the mini-batch (Fig. 1b), which we term batch mode. This feature provides a faster way to deploy augmentations, improving computational efficiency at the expense of sampling diversity.

## Results and Discussion

To benchmark the performance of EvoAug-TF, we utilized the data and deep learning model from the DeepSTARR study^14^. The prediction task is set up to take as input 249 nucleotide (nt) sequences and predict enhancer activity (measured via STARR-seq^15^) for developmental and housekeeping transcriptional promoters in D. melanogaster S2 cells as a multi-task regression. We systematically trained the DeepSTARR model with different EvoAug-TF augmentations individually and in combinations and compared their performance with the original EvoAug, using the same hyperparameters as in the original study^6^. To ensure the robustness of the results, we conducted five trials for each set of augmentations with different random initializations.

We found that models trained with EvoAug-TF augmentations achieved comparable performance to the original EvoAug-trained models with the same augmentation settings (Fig. 1c and 1d). Notably, EvoAug-TF consistently yielded improved performance compared to the original EvoAug after stage 1 (i.e., before finetuning) with the exception of random noise augmentation. However, the original EvoAug yielded slightly better performance than EvoAug-TF upon finetuning. In most cases, models trained with data augmentations led to improved performance compared to standard training.

In terms of computational costs, we found that running EvoAug-TF exhibited similar training times per epoch as EvoAug (Fig. 1e). Thus, EvoAug-TF’s approach of applying the same numbers and types of augmentations to each sequence maintains a comparable computational cost as EvoAug’s approach of applying different numbers and types of augmentations to each sequence. Notably, both approaches are effective in improving model performance.

To further demonstrate the breadth of the EvoAug-TF package, we explored the use of batch mode augmentations. First, we trained convolutional neural networks on ChIP-seq peak classification from the original EvoAug study using batch mode augmentations (i.e., insertion, deletion, and translocation only). As expected, models trained with EvoAug-TF augmentations in batch mode consistently found improved performance (Fig. 1f). We also explored the effectiveness of EvoAug-TF in the low data regime with the same batch mode augmentations. Strikingly, a DeepSTARR model trained with EvoAug-TF on only 25% of the training data yields better performance than the same DeepSTARR model trained on the whole dataset with standard training (Fig. 1g). Thus, EvoAug-TF can greatly improve data efficiency when training genomic deep learning models, and batch mode is an effective way to improve performance over standard training.

In the original study, EvoAug-trained models were found to improve motif representations in attribution maps. To explore whether EvoAug-TF trained models also improve the interpretability of attribution maps, we generated Saliency Maps^16^ for 500 sequences with the highest observed enhancer activity for the developmental promoter and a different set of 500 sequences for the housekeeping promoter in the DeepSTARR dataset. We then quantified the consistency of essential features across a population of attribution maps using the Kullback-Liebler Divergence (KLD) of the distribution of locally embedded attribution scores versus an uninformative prior, which is termed *k*-attr-mean metric^17^. A higher KLD suggests that the attribution scores reflect similar recurring patterns across the 500 attribution maps. Indeed, EvoAug-TF-trained DeepSTARR yielded significantly higher KLD scores than standard training (Fig. 1h).

Attribution methods are sensitive to local function properties and thus can yield spurious attribution scores for reasons that are not biological^17^. To test the robustness of attribution maps, we developed a translational robustness test, similar to the robustness test for model predictions introduced in^5^. The assumption is that small shifts to the sequence should not affect the importance of binding sites (when no significant attribution scores are present near the ends of the sequence). Hence, the attribution scores for the binding sites should also shift with the translations while maintaining the same importance levels. In the translational robustness test, the input sequence was randomly translated by up to 30nt in either direction with np.roll, the (corrected) attribution scores^18^ were calculated for the translated sequence, and then the inverse translation was imposed on the attribution maps to align with the wild type attribution map. This process was repeated 20 times for each sequence, which resulted in 20 translated attribution maps realigned to the input sequence. Next, a variation score was calculated using the root-mean-squared error to summarize the variation across the aligned attribution maps with a scalar value. As expected, DeepSTARR trained with EvoAug-TF yields a significantly lower variability score compared to standard training for Saliency Maps^16^, Integrated Gradients^19^, and DeepSHAP^20^ (Fig. i). Thus, EvoAug-TF training has the benefit of resulting in more robust attribution maps for various attribution methods.

Each augmentation has corresponding hyperparameters that can be tuned to optimize performance gains. Previously, EvoAug identified hyperparameters through a simple grid search, focusing on one augmentation at a time. EvoAug-TF now provides Google Colab notebooks that show how to integrate EvoAug-TF with comprehensive hyperparameter searches, such as population-based training^21^ and the asynchronous hyperband algorithm^22^ provided by Ray Tune^23^. These should offer an alternative strategy to help navigate the combinatoric search space of discovering optimal hyperparameters when deploying multiple augmentations.

## Conclusion

EvoAug-TF is a TensorFlow implementation of EvoAug (a PyTorch package) that provides the ability to train genomic DNNs with evolution-inspired data augmentations. Our results demonstrate the effectiveness of EvoAug-TF to improve generalization and model interpretability with attribution methods. We found that models incorporating EvoAug-TF augmentations achieved comparable or improved performance with similar computational efficiency as the original EvoAug models in PyTorch. Similar to EvoAug, EvoAug-TF is extensible – it can easily accommodate new types of custom augmentations within its data augmentation framework. While the current implementation only supports model outputs as scalars in single- or multi-task settings, we plan to extend these capabilities for multivariate predictions in the future to provide data augmentations for quantitative models that output profiles of read coverages^5,12,13^. Overall, EvoAug-TF offers a new tool for TensorFlow users in the genomics research community to improve data efficiency in training genomic deep learning models, leading to improved generalization and interpretability with attribution analysis.

## Author contributions statement

Y.Y. and P.K. conceived the experiments, Y.Y. developed EvoAug-TF, conducted the experiments, and analyzed the results. S.M. developed the attribution translational robustness test and performed the experiments that used the test. Y.Y., S.M., and P.K. wrote and reviewed the manuscript.

## Acknowledgments

The authors would like to thank Ziqi (Amber) Tang and Chandana Rajesh for support in setting up Ray Tune and to Jakub Kaczmarzyk for support in setting up ReadTheDocs documentation. Research reported in this publication was supported in part by the National Institute Of General Medical Sciences of the National Institutes of Health under Award Number R01GM149921 and the National Human Genome Research Institute of the National Institutes of Health under Award Number R01HG012131. This work was performed with assistance from the US National Institutes of Health Grant S10OD028632-01.

## References

1. Eraslan, G., Avsec, Ž., Gagneur, J. & Theis, F. J. Deep learning: new computational modelling techniques for genomics. Nat. Rev. Genet. 20, 389–403 (2019).

2. Koo, P. K. & Ploenzke, M. Deep learning for inferring transcription factor binding sites. Curr. Opin. Syst. Biol. 19, 16–23 (2020).

3. Zhang, C., Bengio, S., Hardt, M., Recht, B. & Vinyals, O. Understanding deep learning (still) requires rethinking generalization. Commun. ACM 64, 107–115 (2021).

4. Avsec, Ž. et al. Effective gene expression prediction from sequence by integrating long-range interactions. Nat. Methods 18, 1196–1203 (2021).

5. Toneyan, S., Tang, Z. & Koo, P. K. Evaluating deep learning for predicting epigenomic profiles. Nat. Mach. Intell. 4, 1088–1100 (2022).

6. Lee, N. K., Tang, Z., Toneyan, S. & Koo, P. K. Evoaug: improving generalization and interpretability of genomic deep neural networks with evolution-inspired data augmentations. Genome Biol. 24, 105 (2023).

7. Duncan, A. G., Mitchell, J. A. & Moses, A. M. Improving the performance of supervised deep learning for regulatory genomics using phylogenetic augmentation. bioRxiv 2023–09 (2023).

8. Lu, A. X., Lu, A. X. & Moses, A. Evolution is all you need: phylogenetic augmentation for contrastive learning. arXiv 2012.13475 (2020).

9. Lu, A. X. et al. Discovering molecular features of intrinsically disordered regions by using evolution for contrastive learning. PLOS Comput. Biol. 18, e1010238 (2022).

10. Paszke, A. et al. Pytorch: An imperative style, high-performance deep learning library. In Advances in Neural Information Processing Systems 32, 8024–8035 (2019).

11. TensorFlow: Large-scale machine learning on heterogeneous systems.

12. Kelley, D. R. et al. Sequential regulatory activity prediction across chromosomes with convolutional neural networks. Genome research 28, 739–750 (2018).

13. Avsec, Ž. et al. Base-resolution models of transcription-factor binding reveal soft motif syntax. Nat. Genet. 53, 354–366 (2021).

14. de Almeida, B. P., Reiter, F., Pagani, M. & Stark, A. Deepstarr predicts enhancer activity from dna sequence and enables the de novo design of synthetic enhancers. Nat. Genet. 54, 613–624 (2022).

15. Arnold, C. D. et al. Genome-wide quantitative enhancer activity maps identified by starr-seq. Science 339, 1074–1077 (2013).

16. Simonyan, K., Vedaldi, A. & Zisserman, A. Deep inside convolutional networks: Visualising image classification models and saliency maps. arXiv 1312.6034 (2013).

17. Majdandzic, A. et al. Selecting deep neural networks that yield consistent attribution-based interpretations for genomics. In Machine Learning in Computational Biology, 131–149 (PMLR, 2022).

18. Majdandzic, A., Rajesh, C. & Koo, P. K. Correcting gradient-based interpretations of deep neural networks for genomics. Genome Biol. 24, 1–13 (2023).

19. Sundararajan, M., Taly, A. & Yan, Q. Axiomatic attribution for deep networks. In International conference on machine learning, 3319–3328 (PMLR, 2017).

20. Lundberg, S. M. & Lee, S.-I. A unified approach to interpreting model predictions. Adv. Neural Inf. Process. Syst. 30 (2017).

21. Jaderberg, M. et al. Population based training of neural networks. arXiv 1711.09846 (2017).

22. Li, L. et al. Massively parallel hyperparameter tuning. arXiv 1810.05934 (2018).

23. Liaw, R. et al. Tune: A research platform for distributed model selection and training. arXiv 1807.05118 (2018).

